## Supplementary File 1 for "Transcranial direct current stimulation modulates primate brain dynamics across states of consciousness"

### Monkey R. (8 MRI sessions)

#### Anodal electrode (prefrontal)

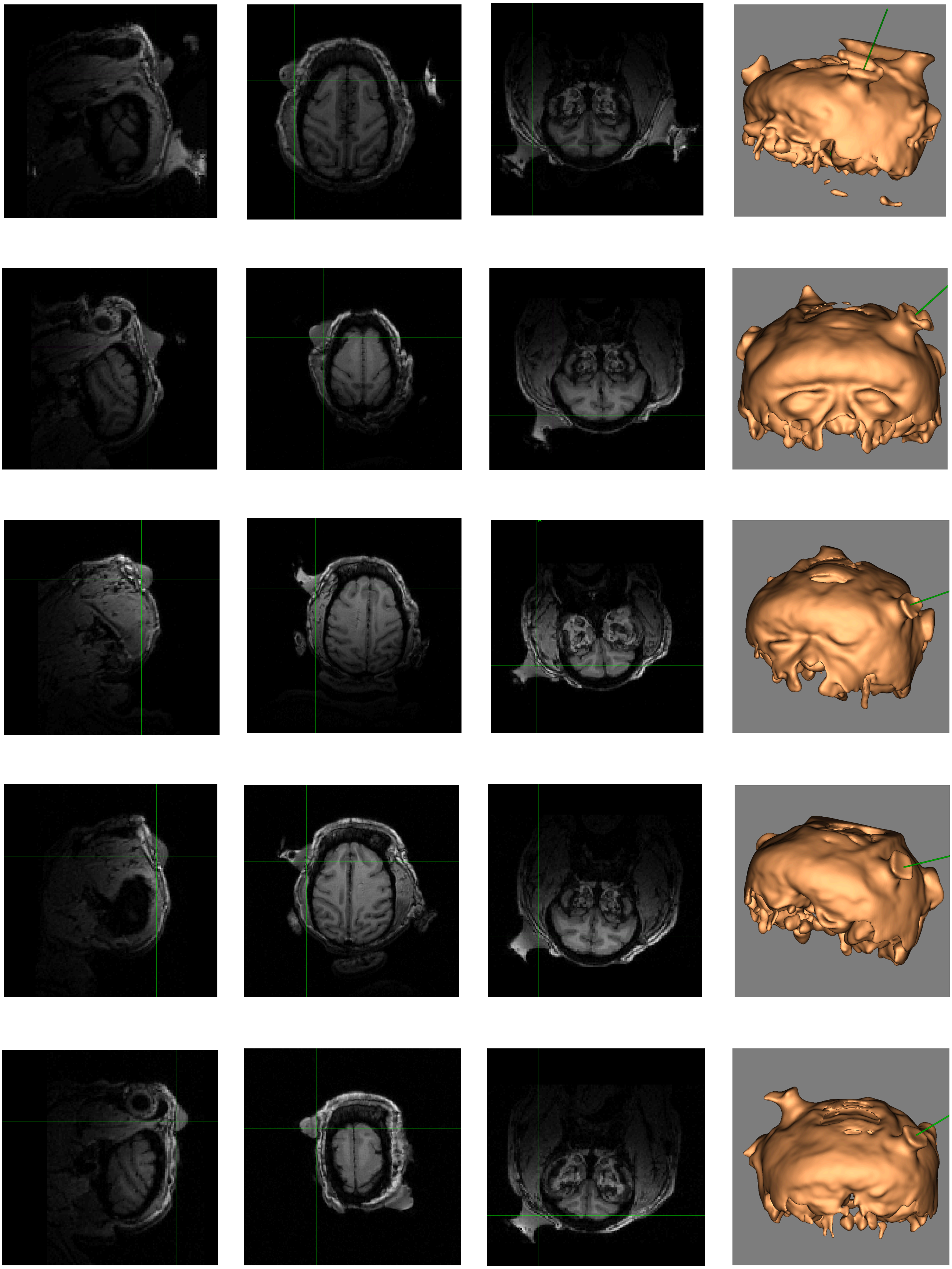

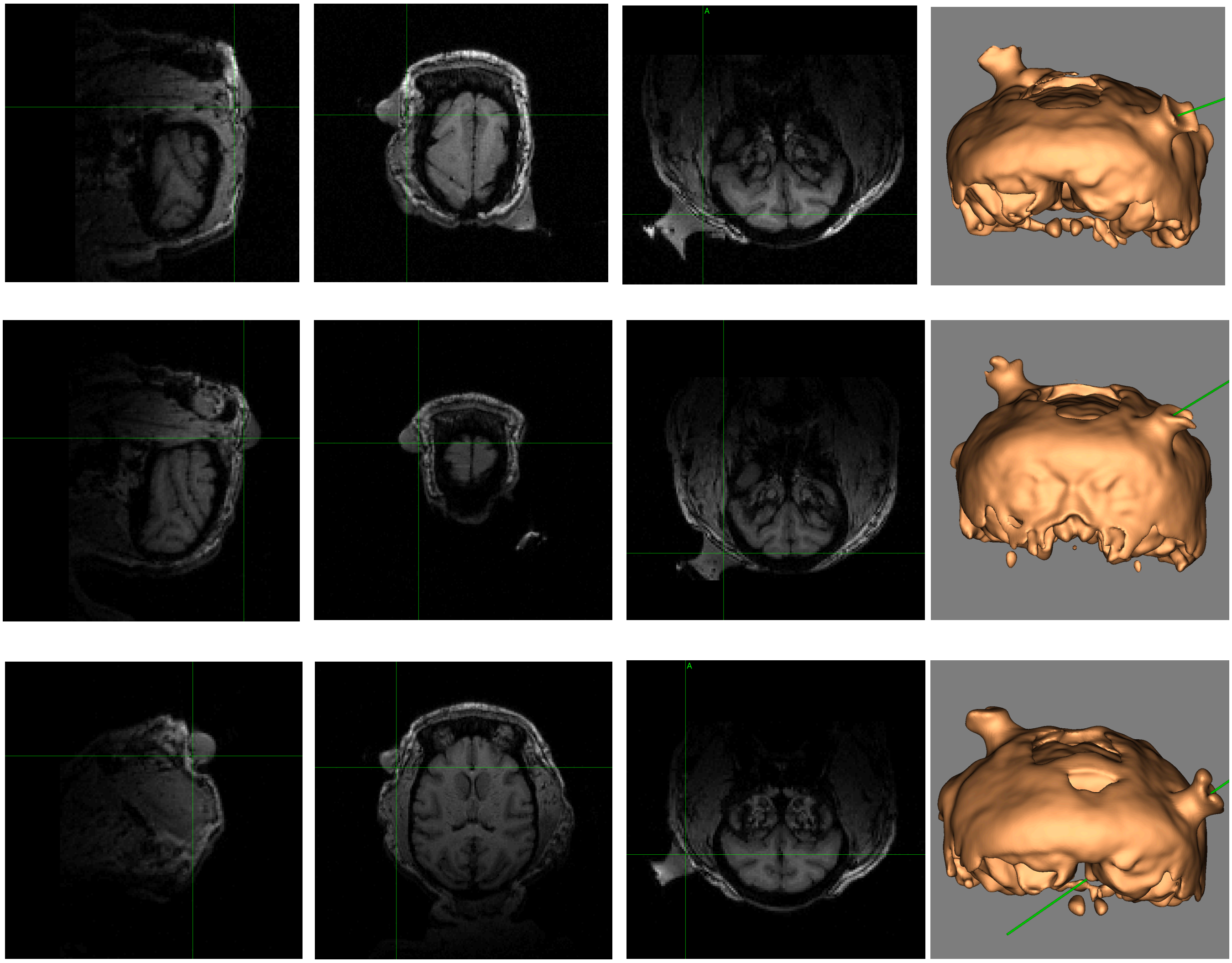

#### Cathodal electrode (occipital)

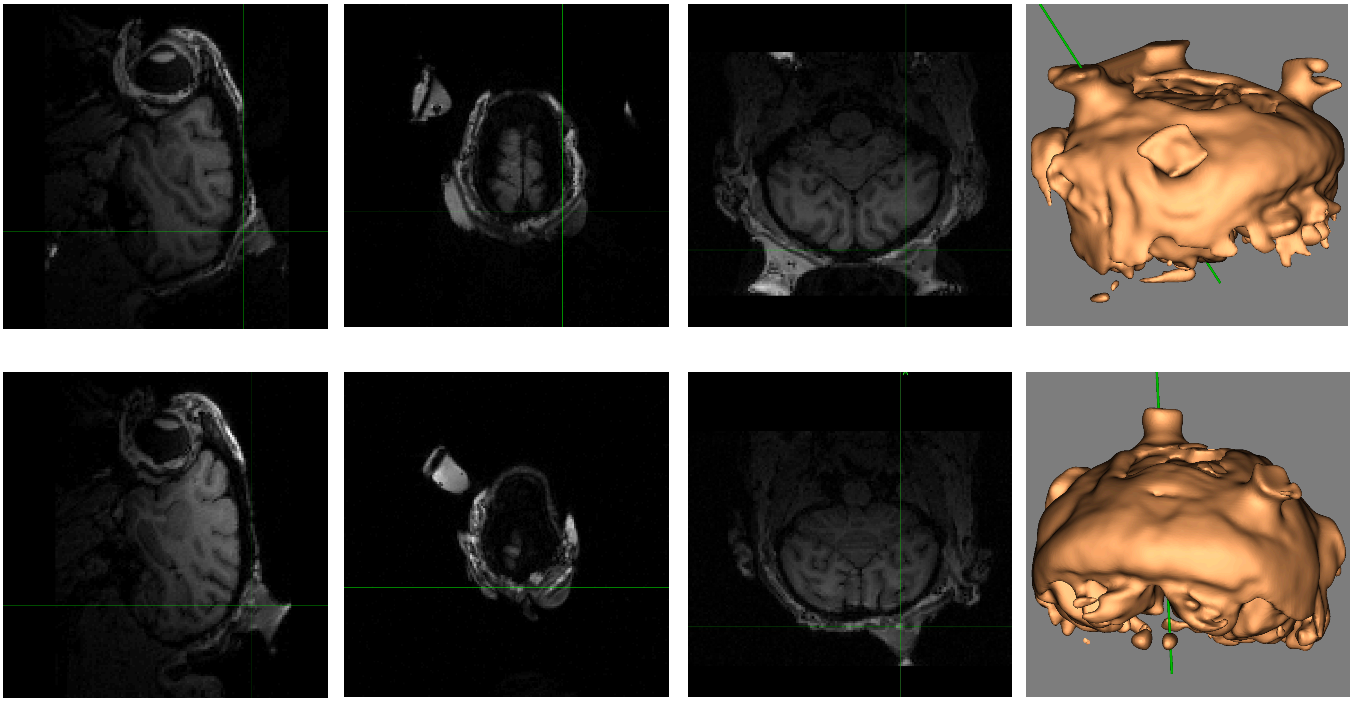

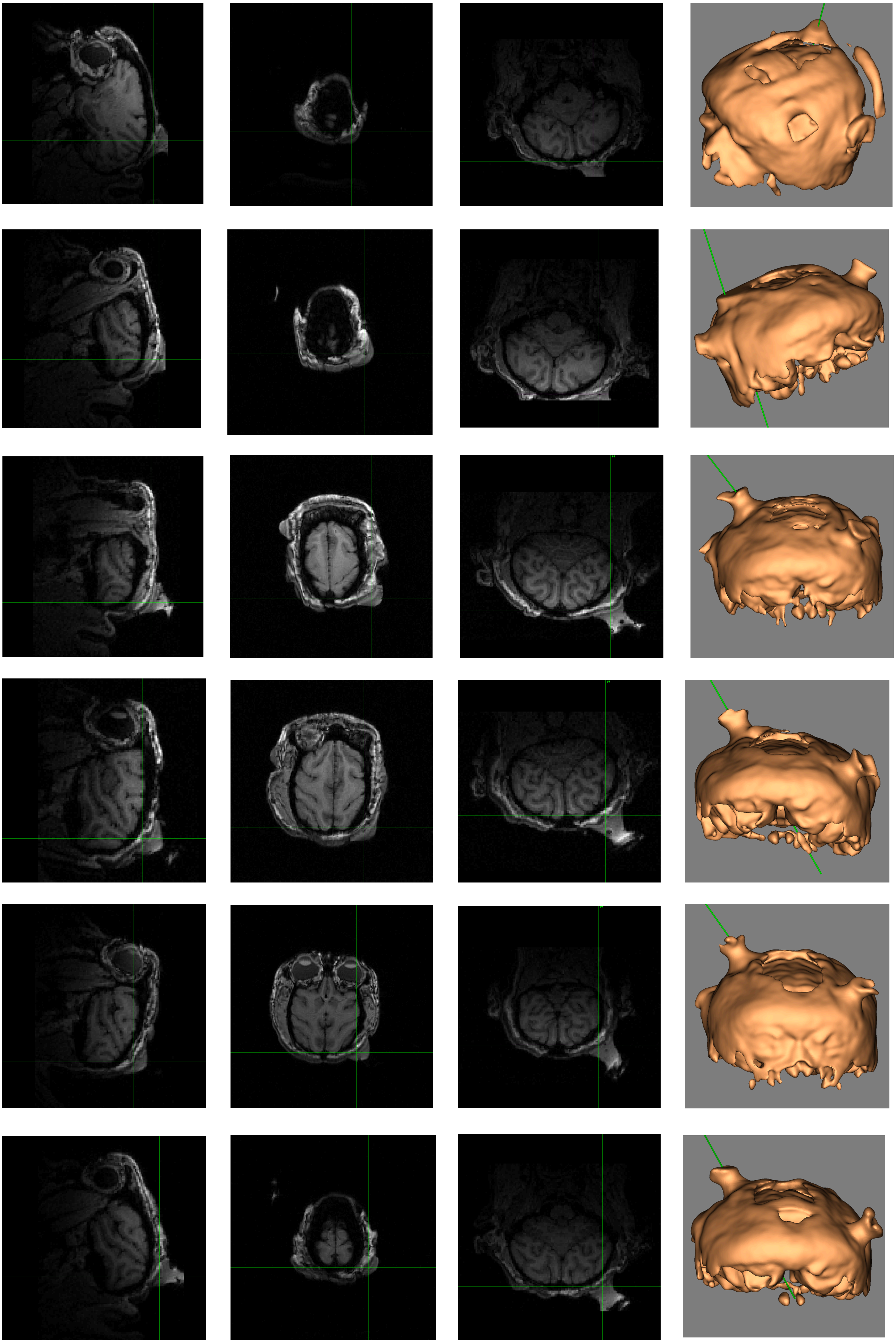

### Monkey N. (2 MRI sessions)

#### Anodal electrode (prefrontal)

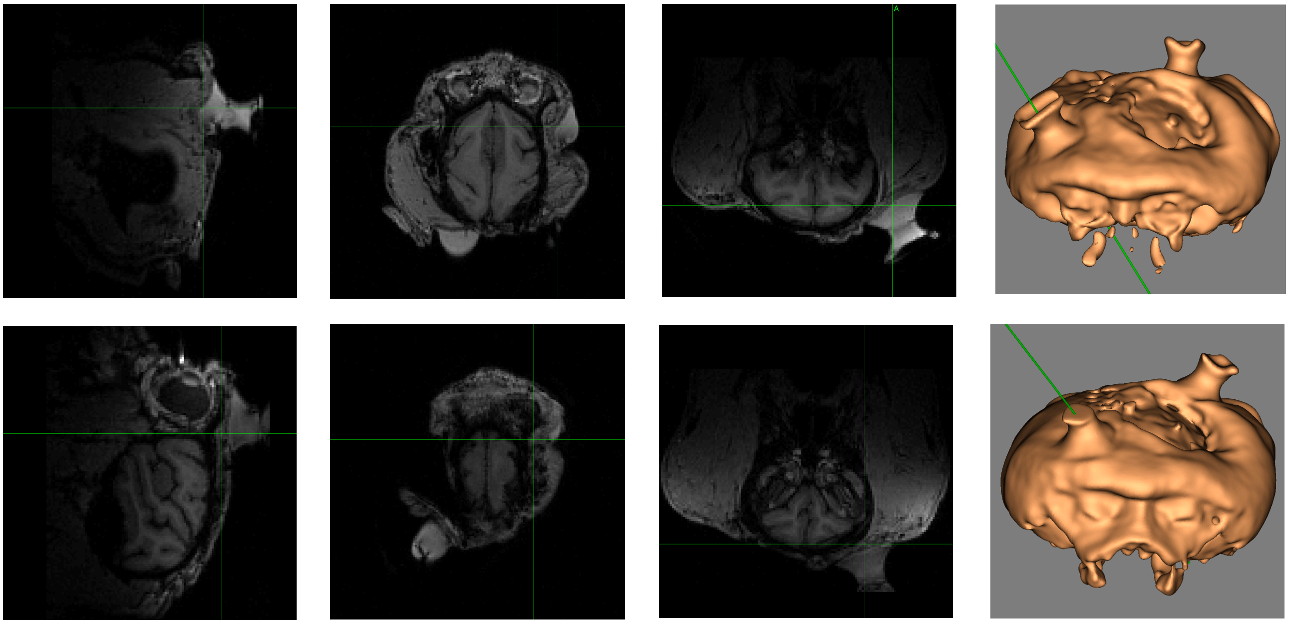

#### Cathodal electrode (occipital)

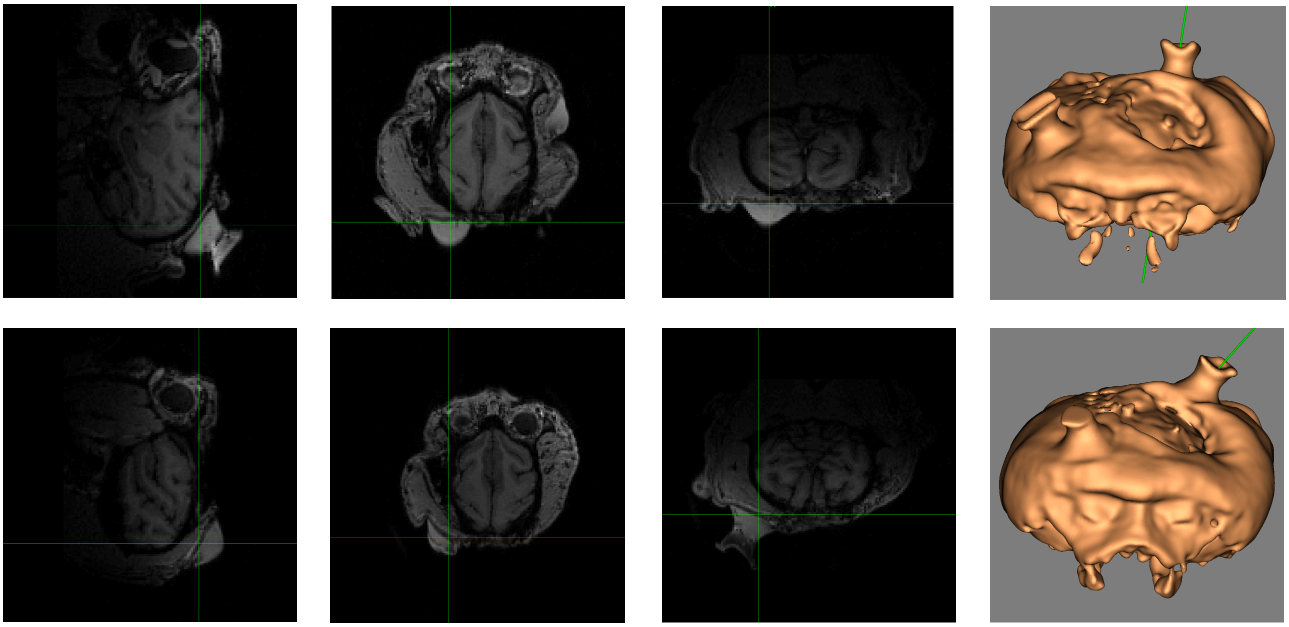

### Monkey J. (16 MRI sessions)

#### Anodal electrode (prefrontal)

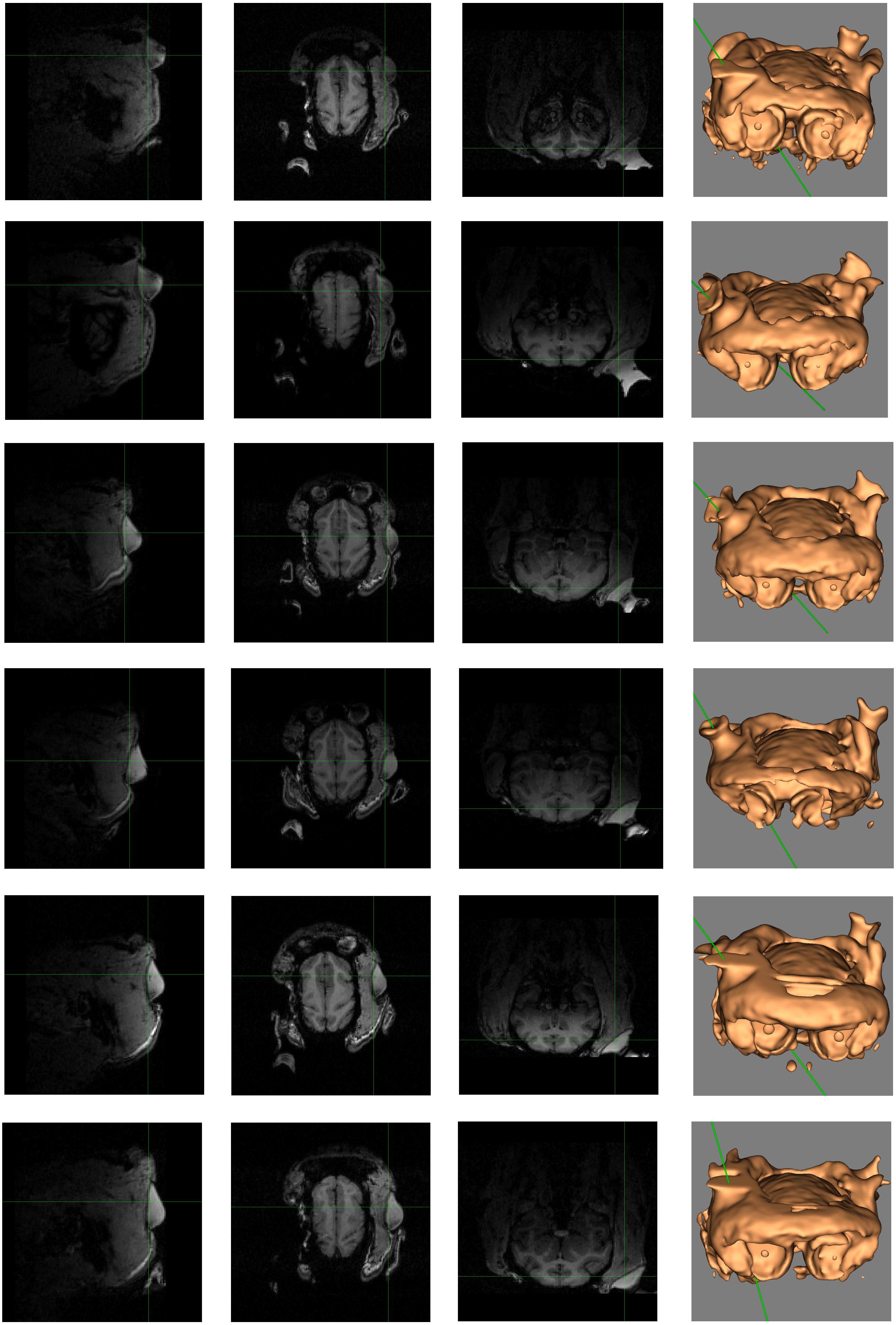

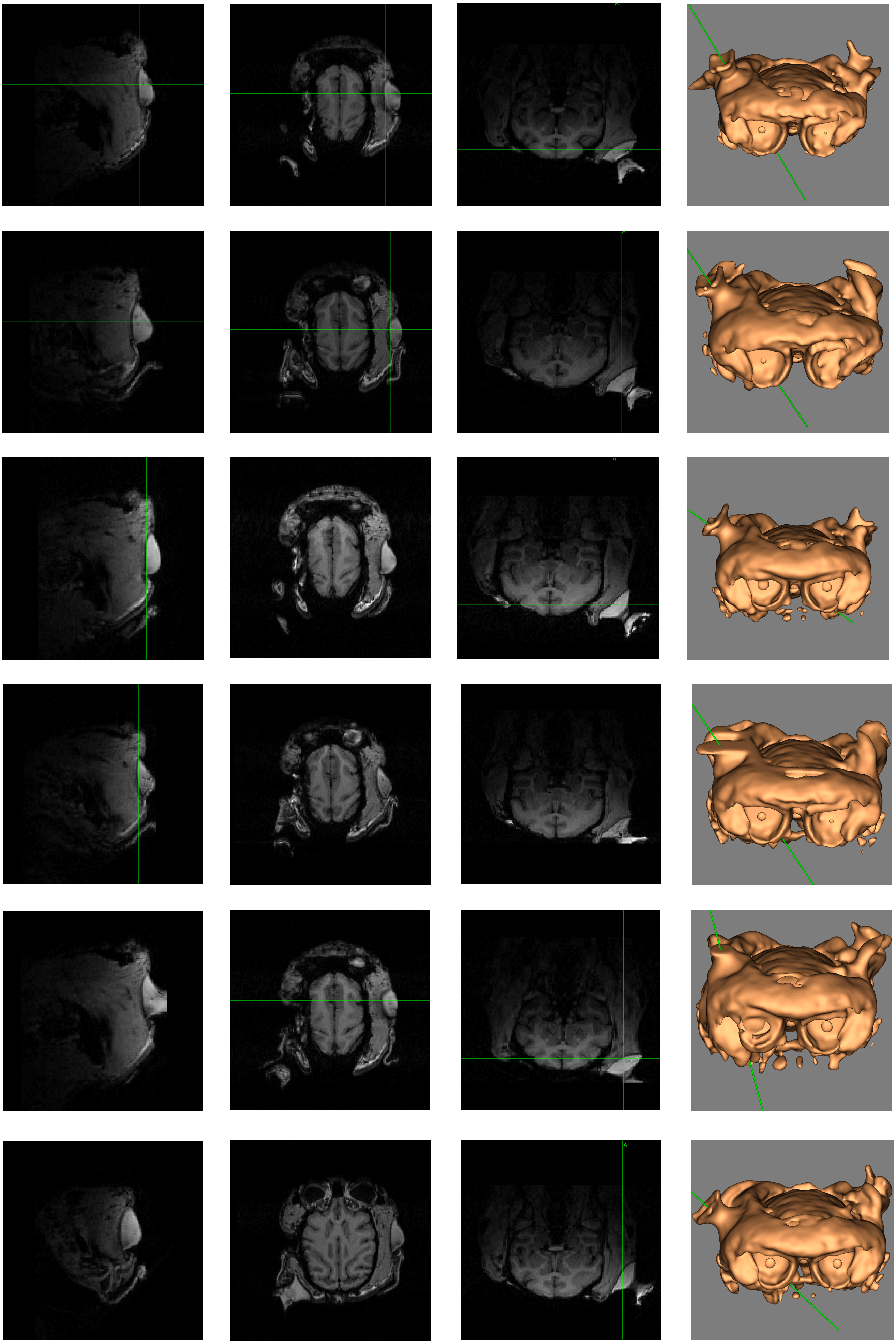

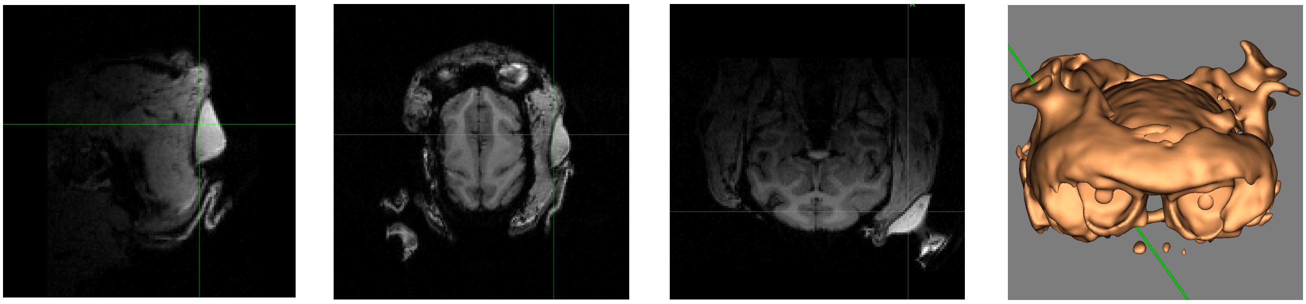

#### Cathodal electrode (prefrontal)

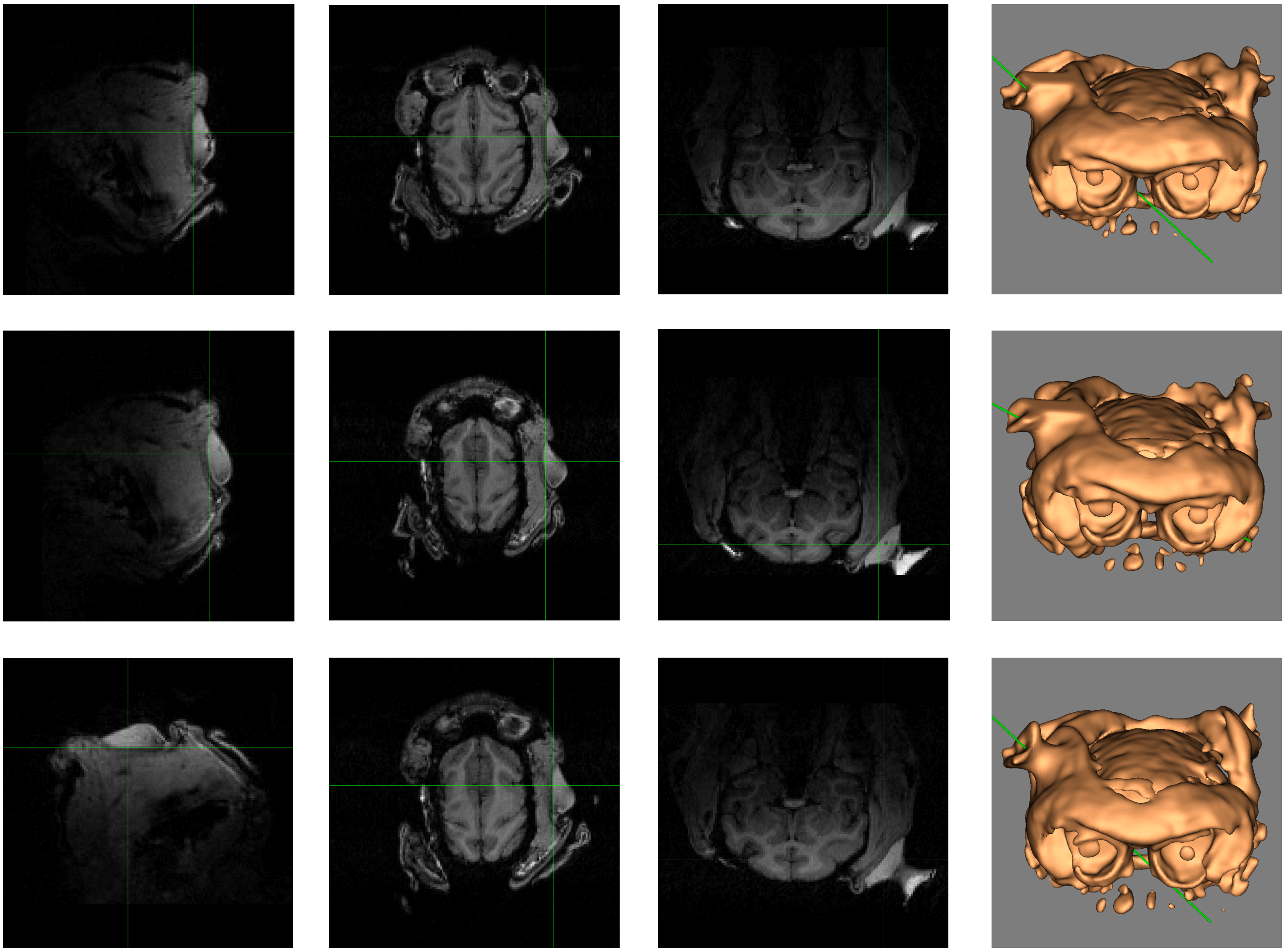

### Cathodal electrode (occipital)

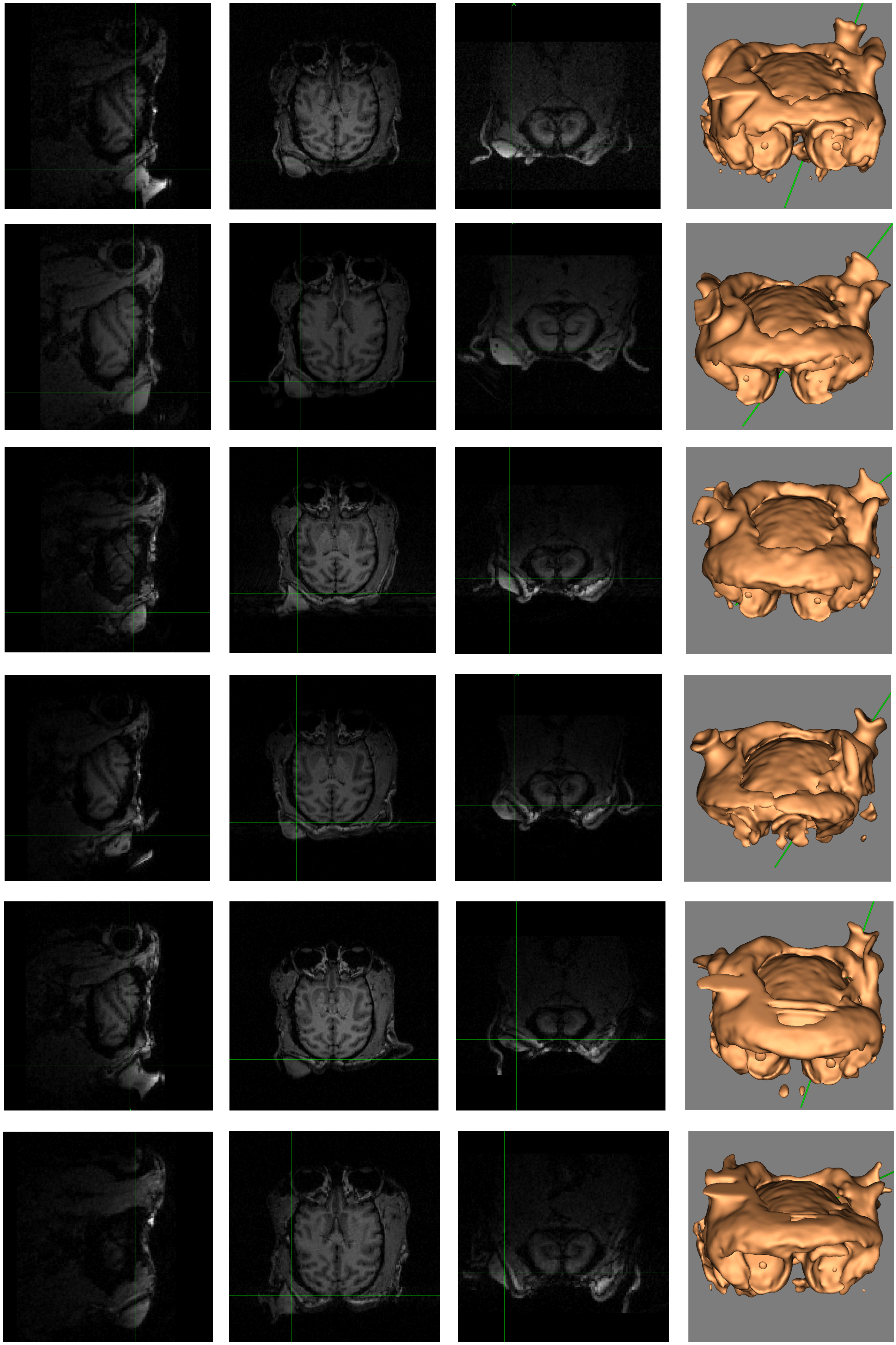

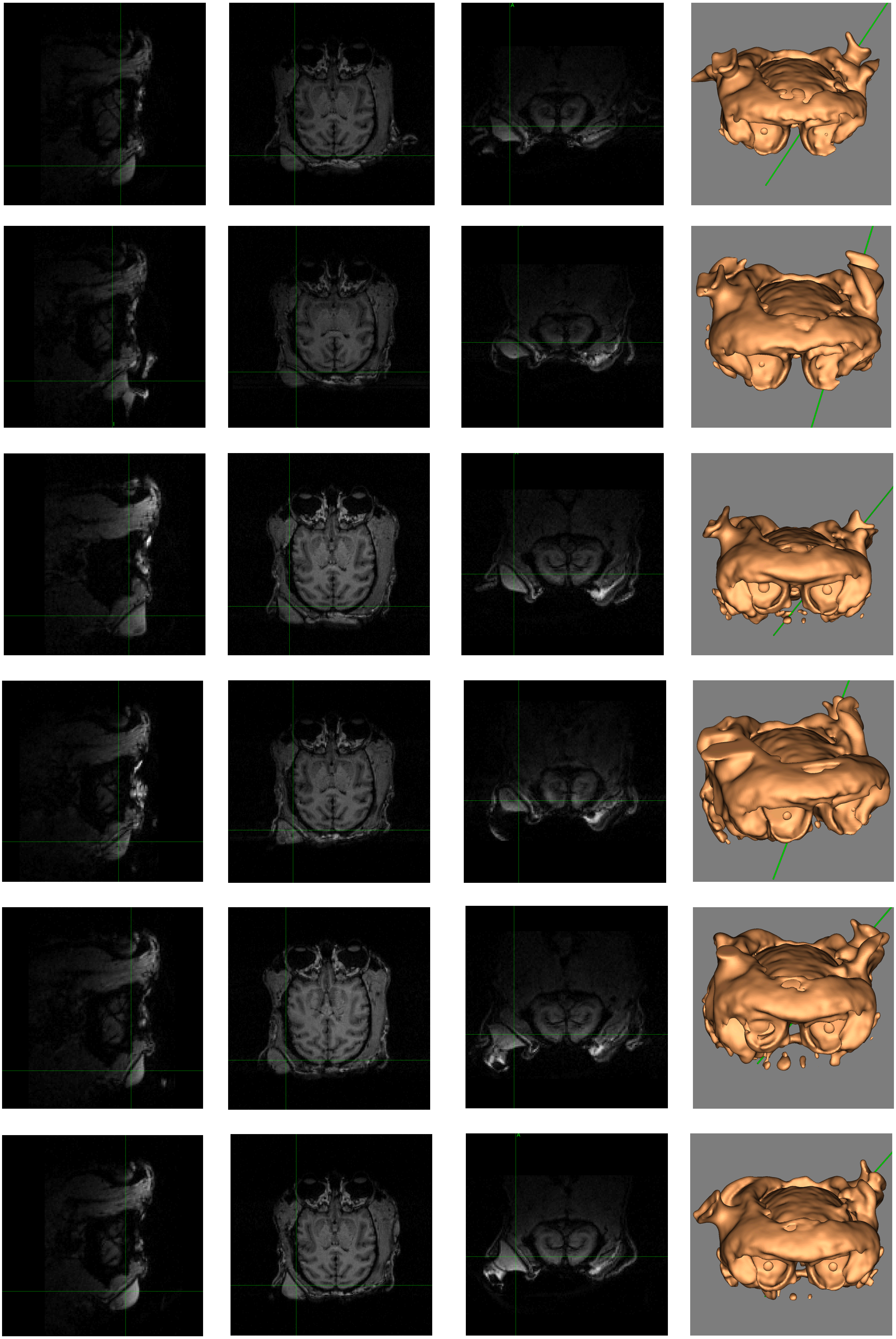

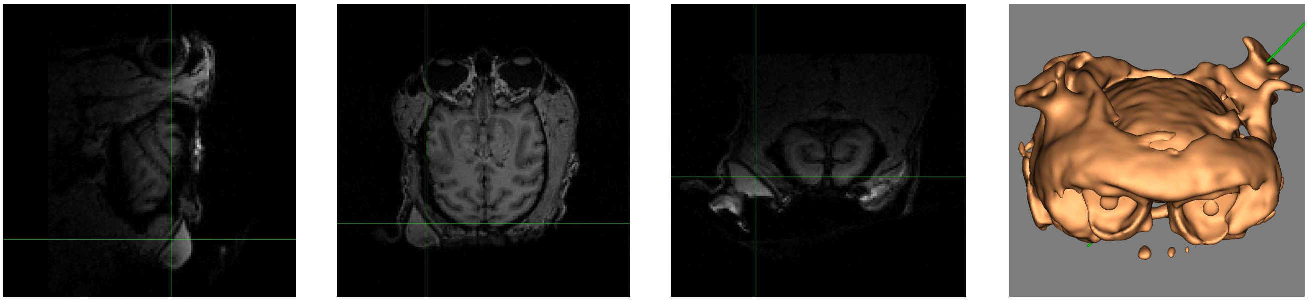

#### Anodal electrode (occipital)

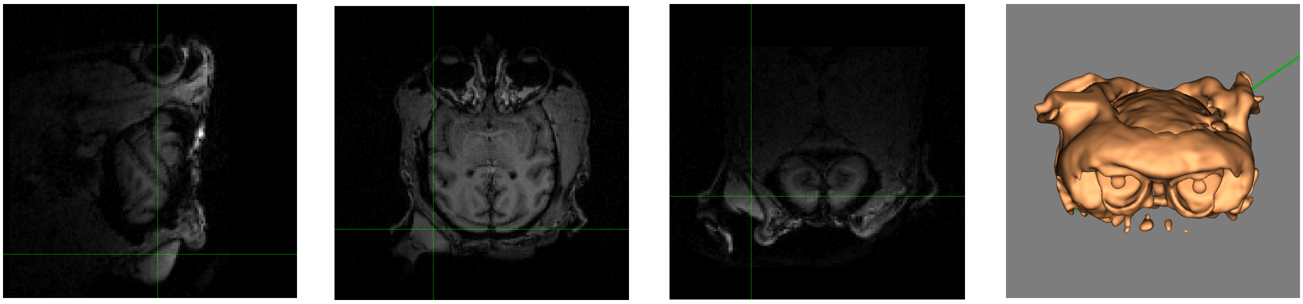

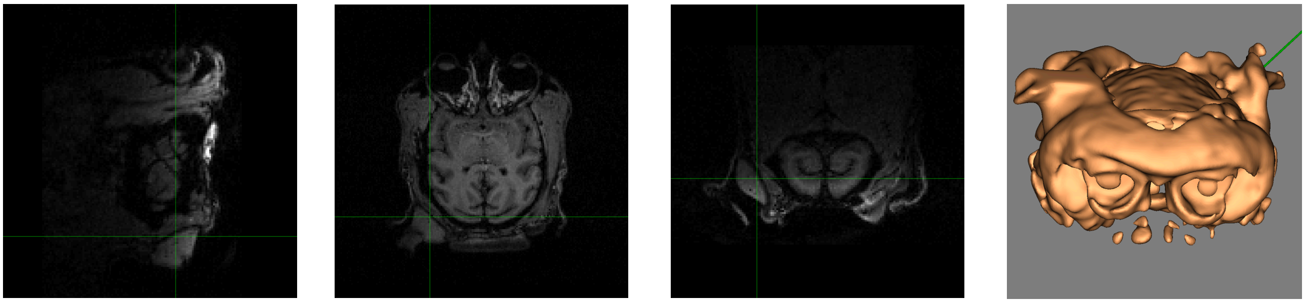

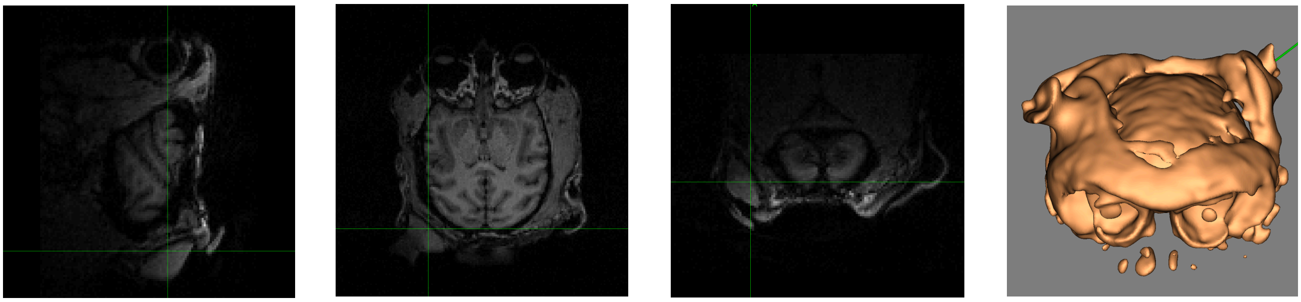

### Monkey Y. (14 MRI sessions)

#### Anodal electrode (prefrontal)

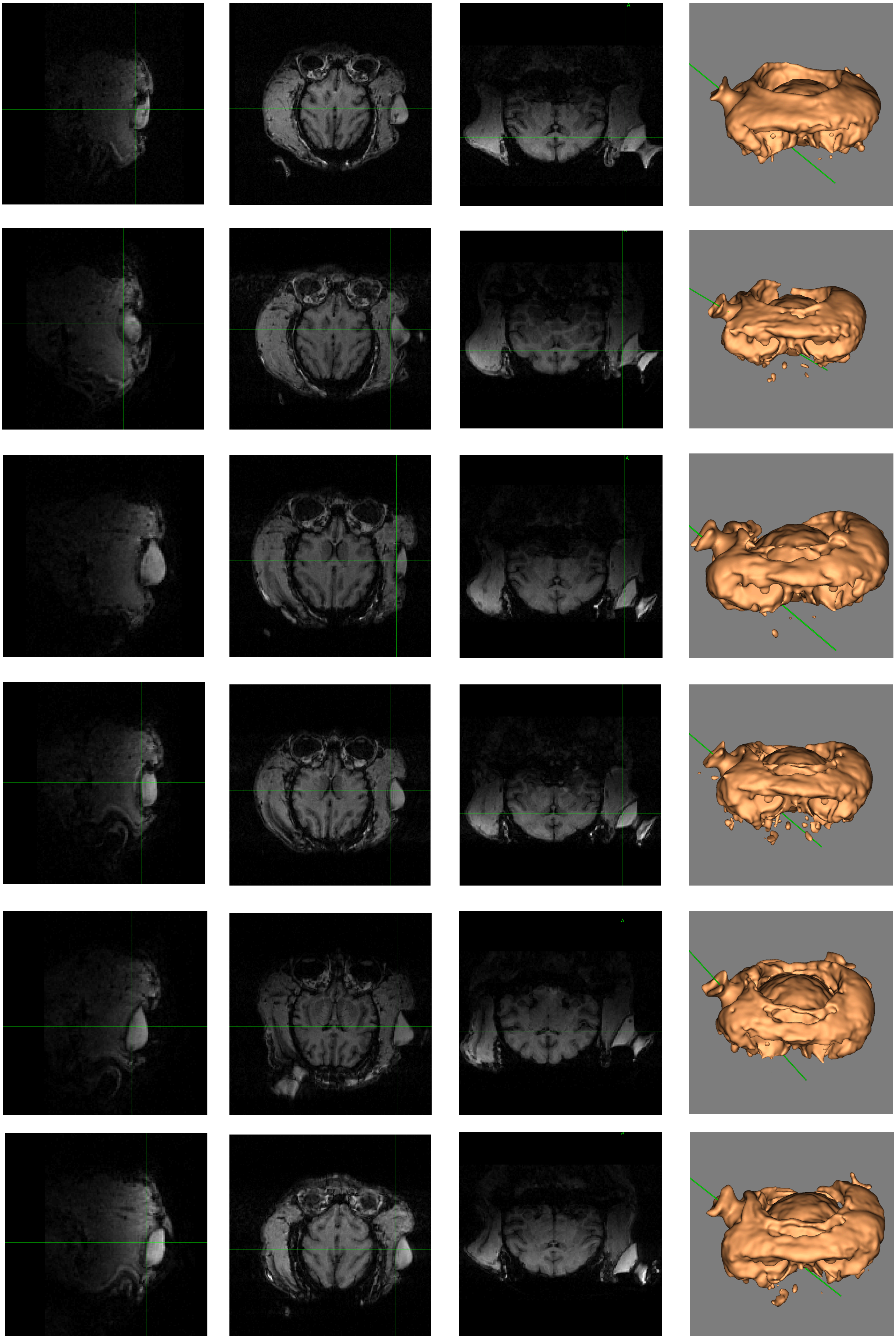

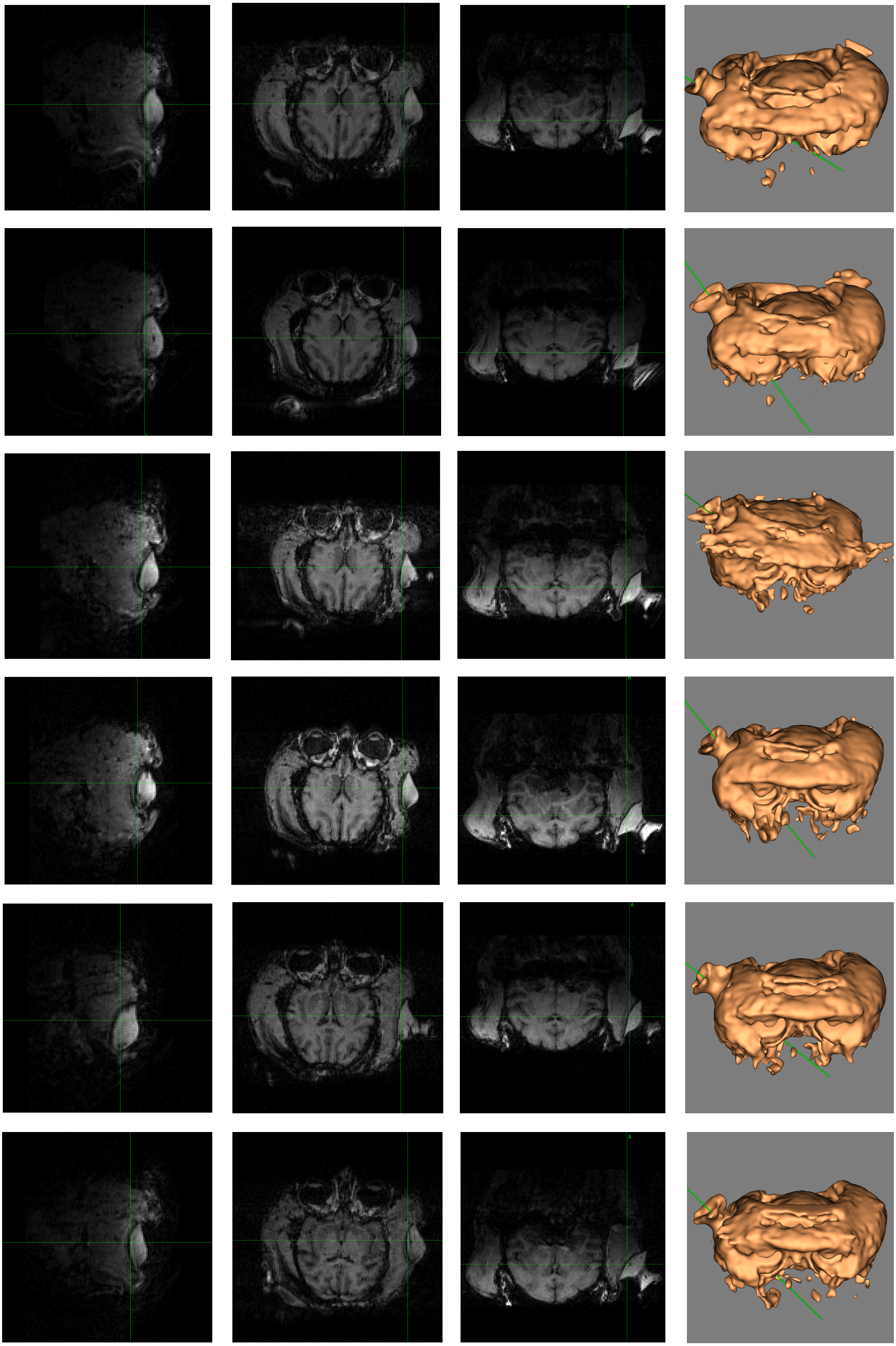

#### Cathodal electrode (prefrontal)

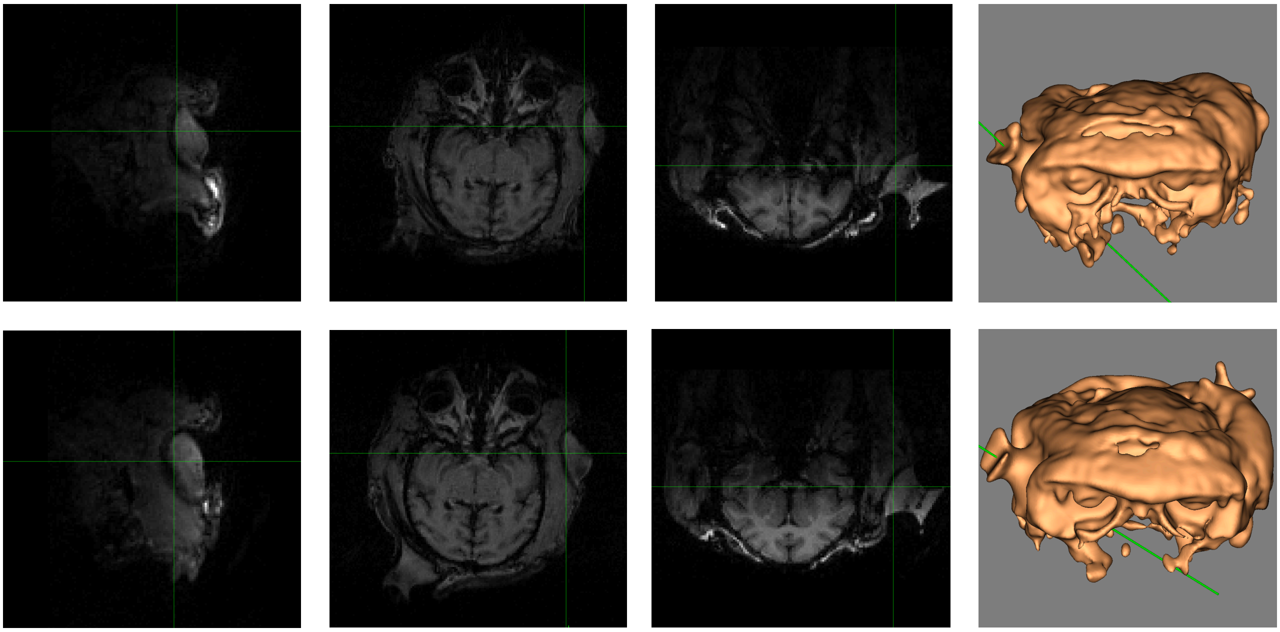

#### Cathodal electrode (occipital)

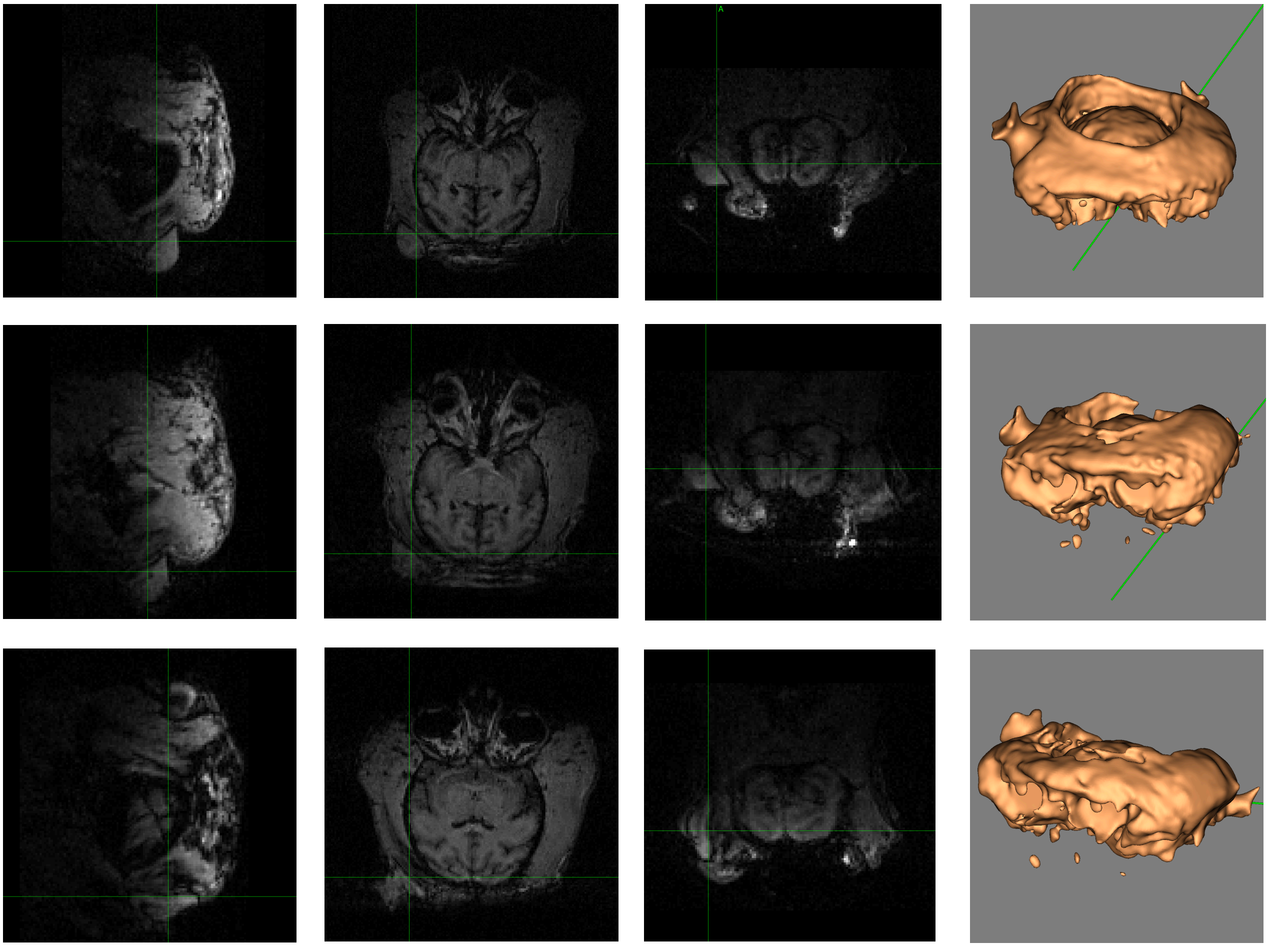

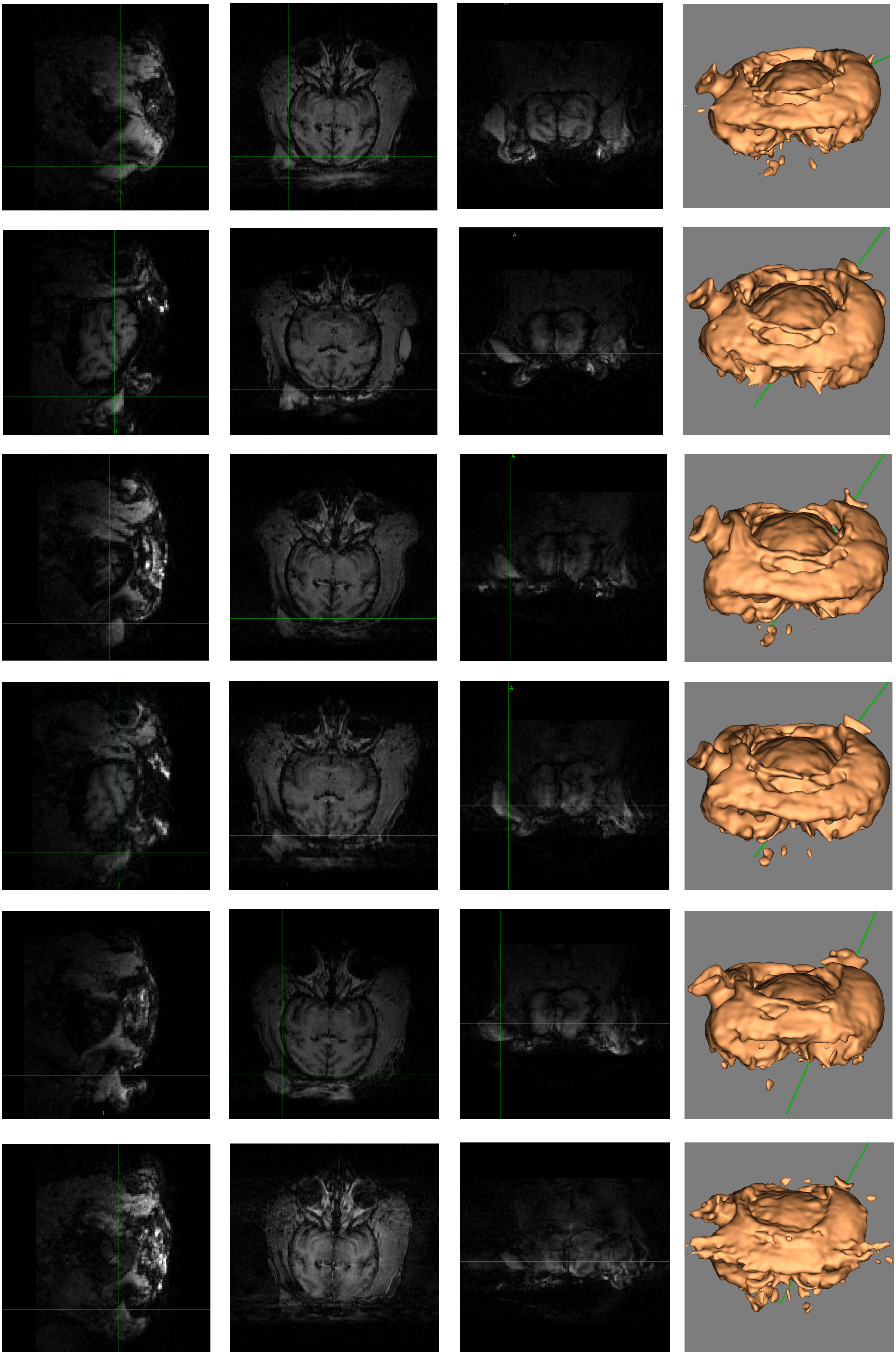

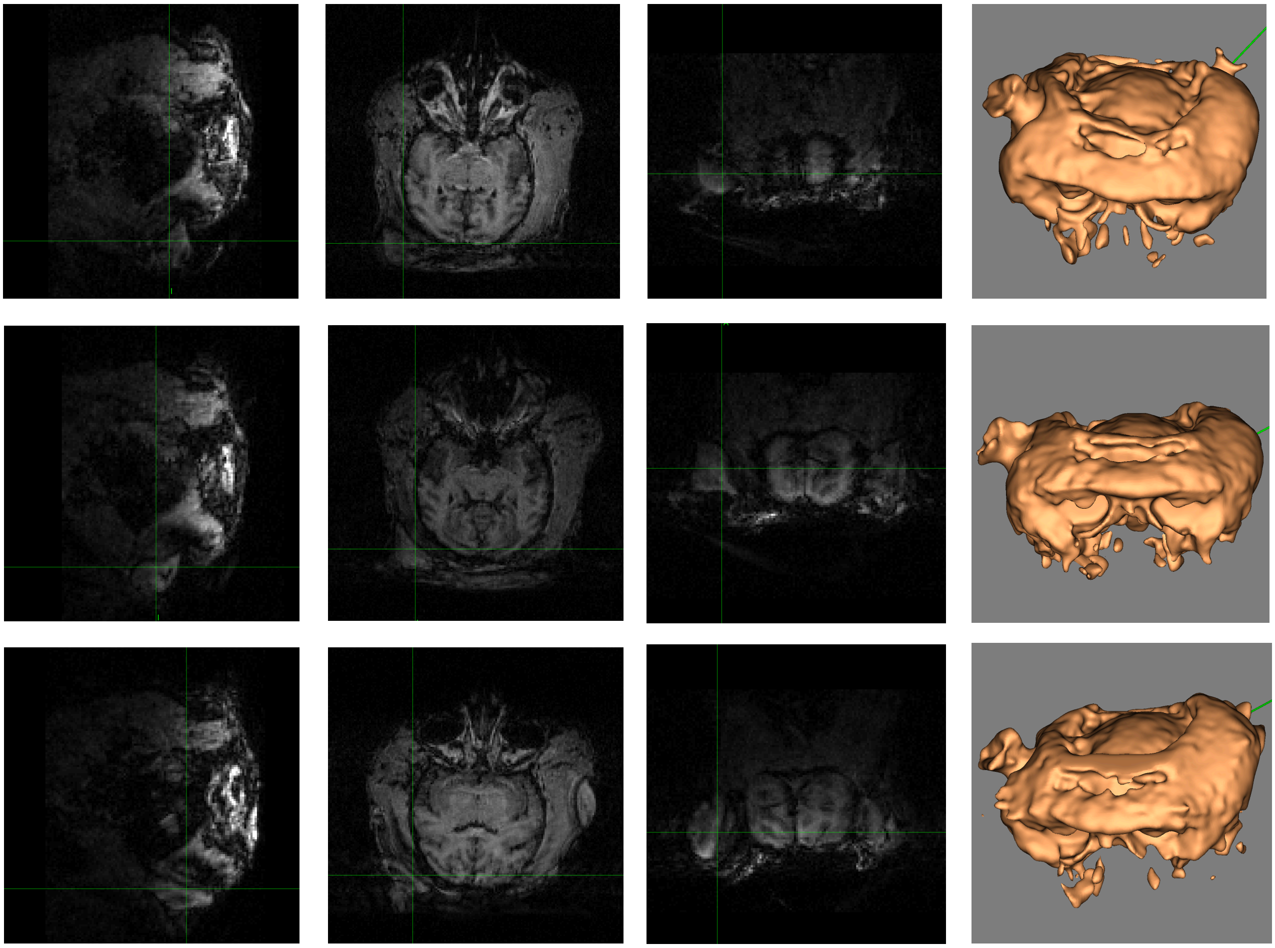

#### Anodal electrode (occipital)

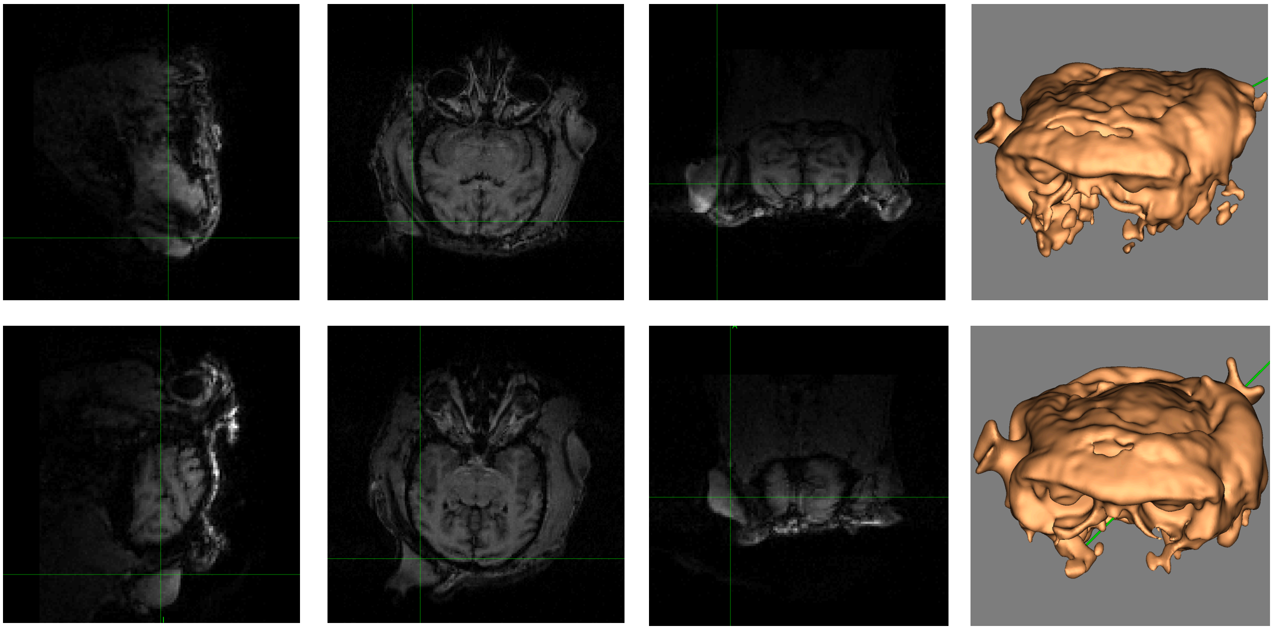
