## Supplementary figures and images for "Transcranial direct current stimulation modulates primate brain dynamics across states of consciousness"

### Supplementary File 2

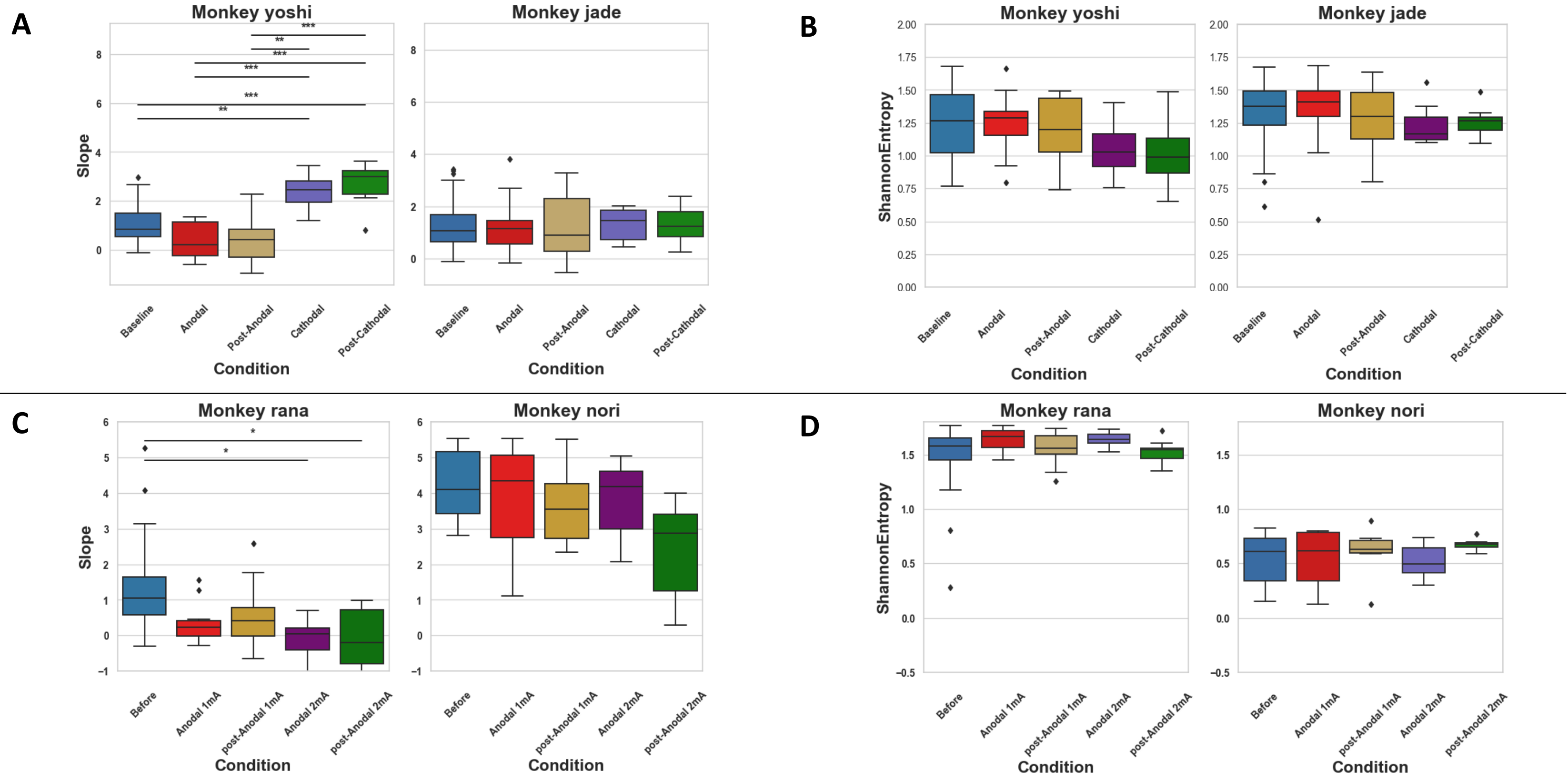

### Supplementary File 3

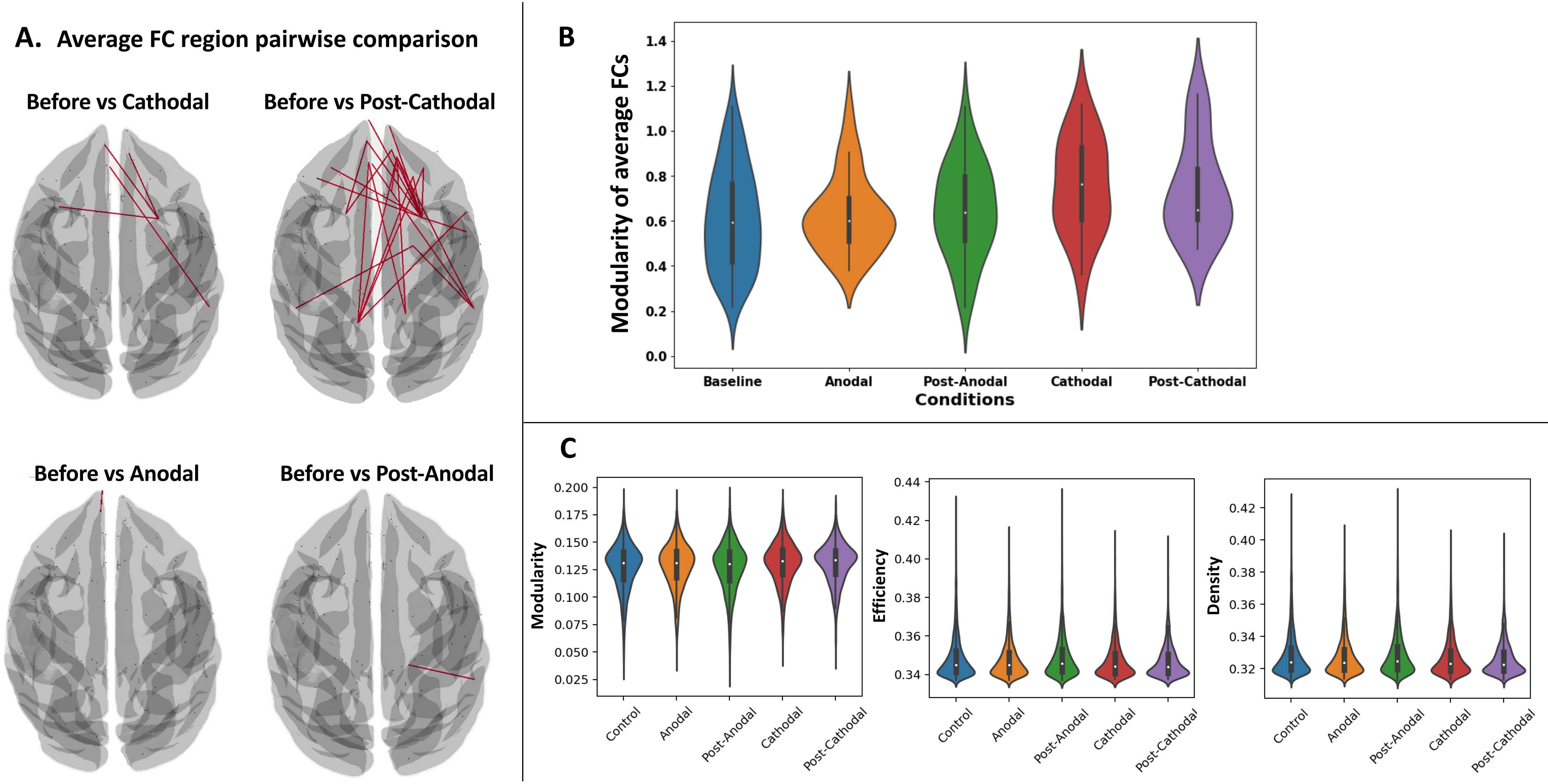

### Supplementary File 4

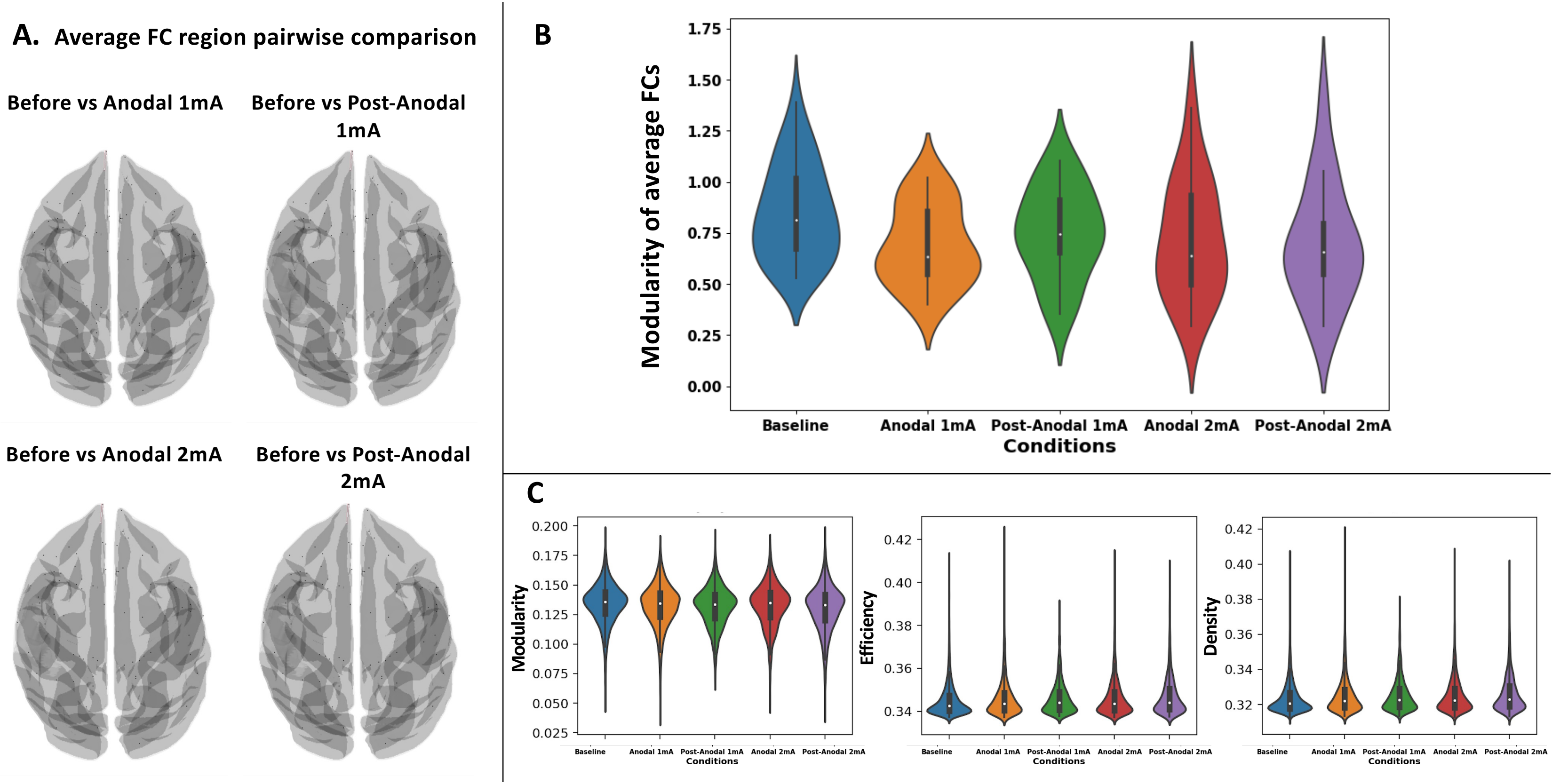
